# Explainable t-SNE for single-cell RNA-seq data analysis

**DOI:** 10.1101/2022.01.12.476084

**Authors:** Henry Han, Tianyu Zhang, Chun Li, Mary Lauren Benton, Juan Wang, Junyi Li

## Abstract

**Background:** Single-cell RNA (scRNA-seq) sequencing technologies trigger the study of individual cell gene expression and reveal the diversity within cell populations. To measure cell-to-cell similarity based on their transcription and gene expression, many dimension reduction methods are employed to retrieve corresponding low-dimensional embeddings of input scRNA-seq data to conduct clustering. However, the methods lack explainability and may not perform well with scRNA-seq data because they are not customized for high-dimensional sparse scRNA-seq data.

**Results:** In this study, we propose an explainable t-SNE: cell-driven t-SNE (c-TSNE) that fuses cell differences reflected from biologically meaningful distance metrics for input data. Our study shows that the proposed method not only enhances the interpretation of the original t-SNE visualization but also demonstrates favorable single cell segregation performance on benchmark datasets compared to state-of-the-art peers. The robustness analysis shows that the proposed cell-driven t-SNE demonstrates robustness to dropout and noise in clustering. It provides a novel and practical way to investigate the interpretability of t-SNE in scRNA-seq data analysis. Unlike the general assumption that the explainability of a machine learning method may need to compromise with learning efficiency, the proposed explainable t-SNE improves both clustering efficiency and explainability. More importantly, our work suggests that widely used t-SNE can be easily misused in existing scRNA-seq analysis, because its default Euclidean distance can bring biases or meaningless results in cell difference evaluation for high-dimensional sparse scRNA-seq data. To the best of our knowledge, it is the first explainable t-SNE proposed in scRNA-seq analysis and will inspire other explainable machine learning method development in the field.

**Conclusion:** The proposed explainable t-SNE outperforms classic t-SNE and its peers in meaningful visualization and segregation. The poor performance of the classic t-SNE highlights the importance of developing explainable machine learning methods in scRNA-seq analysis. The explainable t-SNE is a data-centric customized ML enhance efficiency in data analysis through bringing more biological insights and interpretations.

## Background

The recent emergence of the single-cell RNA sequencing (scRNA-seq) enables the study of gene expression at the level of individual cells, which can give more insight into the unexplained heterogeneity among cell populations [1]. scRNA-seq unveils transcriptomic landscapes by detecting the high-resolution differences between cells. Compared to bulk RNA-seq that assesses bulk cell populations, it measures gene expression on the level of individual cells, and deciphers essential cell-to-cell variability, and uncovers the dynamics of cell fate decision making by precisely comparing the transcriptome of individual cells. It brings more insight into the unexplained heterogeneity among cell populations by unveiling latent subtle biological behaviours compared to bulk RNA-seq and other traditional profiling techniques. The revelation of heterogeneity across each cell contributes to the identification of new cell subpopulation, which is particularly important in unveiling the mechanism of complex diseases, immune systems, and neural systems [2].

However, to reveal the heterogeneity among cell populations, it is essential to quantify variations between each cell’s gene expression profile via clustering scRNA-seq phenotypes. In other words, the critical question that must be answered is which cells are similar or different based on their transcription and gene expression via clustering. scRNA-seq clustering (segregation) remains a challenge in machine learning and bioinformatics because scRNA-seq data is high-dimensional sparse data with built-in random noise from sequencing and experimental design artifacts. Its number of variables is much larger than that of observations. Each observation (sample) contains a large number of zeros or near-zero values because of dropouts, where expressed transcripts may not be detected and assigned zero expression values in sequencing [2]. The noise can be rooted in batch effects, low sequencing depth, dropouts, or biological factors such as the stochastic mechanism in gene expression. Although different methods (e.g., ZIFA) have been developed to deal with the zero-inflation problem, it remains unclear how effectively they can enhance scRNA-seq clustering compared to other methods [3]. Therefore, the high-dimensionality, high-nonlinearity, and zero-inflation along with noise present a hurdle for effective scRNA-seq clustering, especially because existing normalization methods are still premature in most studies.

Many scRNA-seq studies have employed state-of-the-art dimension reduction algorithms such as t-SNE (t-distributed stochastic neighbor embedding) in scRNA-seq clustering [4–6]. They map scRNA-seq samples in an alternative low-dimensional space to detect subtle differences across samples and seek subpopulation similarities [7]. Although exceptions exist, the dimension reduction techniques provide a transformed feature extraction procedure to de-noise data, reduce redundancy between variables, and visualize data from different perspectives to unveil latent data behaviors. For example, t-SNE seeks a low-dimensional embedding for input data to keep the original intrinsic structure by maintaining the relative entropy between the probability distributions induced by the pairwise distances in the input and low-dimensional space [8]. Compared to the holistic dimension reduction methods such as PCA, t-SNE and UMAP (uniform manifold approximation and projection) are good at capturing subtle local data behaviors. Sun et al provided detailed evaluations and comparisons about different dimension reduction methods on various scRNA-seq datasets in scRNA-seq clustering [4].

However, most methods do not perform as well as expected in scRNA-seq clustering. Many researchers believe the special characteristics of scRNA-seq data play an important role in the issue, because scRNA-seq data may not match the methods well [8]. Another important but rarely mentioned factor is that the dimension reduction methods employed are not explainable. They act as a black-box for users rather than provide understandable data-driven interpretations. For example, t-SNE is a widely employed dimension reduction method in scRNA-seq clustering with decent performance. However, it can be hard to explain why there are almost no methodological differences when applying t-SNE to a low-dimensional high-frequency trading dataset versus a high-dimensional sparse scRNA-seq dataset. The former has 10,000+ observations and around 10 variables with almost zero sparseness; The latter has less than 500 observations and more than 20,000 variables, with at least 20% sparseness, in which at least 20 precents of entries are zeros [8].

Although most t-SNE applications employ the default Euclidean distance in t-SNE to compute the required pairwise distance matrix, the Euclidean distance may complicate interpretation or even introduce bias for high-dimensional sparse scRNA-seq data. This is because the Euclidean distance gives the same importance to each direction of an input sample, which is usually expected to be dense data that has almost zero sparseness. It is doubtful that this metric can be applied well to high-dimensional sparse scRNA-seq data without generating biases. Previous work also pointed out the Euclidean distance can be meaningless when applied to high-dimensional data because of ‘curse of high-dimensionality’ as well as a high computing burden [9]. Therefore, the sample similarity calculation should be more customized for scRNA-seq data for the sake of interpretation and accuracy. In summary, though the state-of-the-art t-SNE method is widely used in scRNA-seq clustering and achieves acceptable performance, it is still not an explainable machine learning method for scRNA-seq data because it is simply migrated from machine learning without considering the nuances of the input data.

Although explainable machine learning is still a new topic in biomedical data science, they can be essential for scRNA-seq data analysis. Explainable machine learning will contribute to transparency and accuracy in clustering and other downstream analysis by avoiding biased results, besides enhancing trustworthiness of machine learning results. Furthermore, scRNA-seq data analysis is closely bundled with disease diagnosis. This high-stakes application domain usually has higher expectations for the interpretation of machine learning methods

How can we develop an explainable t-SNE method for the sake of scRNA-seq clustering? The t-SNE embedding is generally used for single cell segregation for its advantage in clustering accuracy and complexity [4,8]. Enhancing the explainability of t-SNE will make t-SNE more applicable to single cell data analysis and provide more accurate and robust cell segregations, but the explainability of t-SNE is rarely investigated in almost all existing scRNA-seq analysis. It is probably because the explainability of dimension reduction is seldom mentioned in the existing machine learning literature because most research efforts are investigated in the explainability of supervised machine learning [10–11].

We believe an explainable t-SNE will be a customized t-SNE that considers the special characteristics of input scRNA-seq data. It should employ more biologically explainable distance metrics to evaluate scRNA-seq sample similarity rather than only use the default. The explainable distance metrics would bring better cell discrimination and contribute to better segregation. Therefore, in contrast with the assumption that there is a trade-off between explainability and learning efficiency in machine learning, the explainable t-SNE should be able to enhance low-dimensional embedding quality for the sake of clustering compared to the original t-SNE because it is customizedly designed according to input data [11–12].

On the other hand, the performance of many dimension-reduction and following clustering algorithms depend critically on the distance metric to calculate sample similarity in the input space. It remains unknown which distance metrics will be appropriate for scRNA-seq data. Wang *et al*. proposed a SIMLR (single-cell interpretation via multi-kernel learning) approach to learn an appropriate similarity metric between samples by employing multi-kernel learning [8]. The learned distance matrix in SIMLR would represent the sample similarities automatically through an optimization framework. Their results show that SIMLR would outperform most of peer methods in scRNA-seq clustering [8]. This seminal work suggests that cell diversity should be evaluated by more advanced distances rather than the default Euclidean distance.

However, SIMLR lacks good interpretability because the results may not be explained well on behalf of scRNA-seq data sample diversity. It is unknown whether the distance metric learned from SIMLR is biologically meaningful. There is also no guarantee that SIMLR will gain an optimal distance metric for sample similarity discrimination via multi-kernel learning [8]. It remains unclear whether the inferred distance would match high-dimensional sparse data well because their method theoretically can be applied to any generic data rather than only designed for scRNA-seq data. Besides, they may not explain well how the multi-kernel functions reveal internal differences between cells’ gene expressions [8]. On the other hand, quite a few imputation methods were proposed for the sake of scRNA-seq clustering from different perspectives (e.g., low-rank approximation) [13–14]. They provide alternative ways to overcome the dropout issue in scRNA-seq analysis. In this investigation, however, we view dropout as a built-in characteristic in sampling transcriptome for each scRNA-seq data sample rather than conduct imputation for clustering. Therefore, our input data is raw single cell RNA-seq data before normalization or imputation.

In this study, we propose an explainable t-SNE method: cell-driven t-SNE (c-TSNE) for scRNA-seq visualization and clustering. It is a customized t-SNE augmented with cell-driven distance metrics. Unlike the classic t-SNE, c-TSNE provides biologically meaningful interpretations to cell discriminations and more explainable low-dimensional embeddings. The customized cell-driven metrics are designed to reflect cell differences between scRNA-seq samples from a biologically explainable viewpoint. They make relevant samples mapped as the closest neighbors to each other in the low-dimensional embedding space in dimension reduction. Different from the general t-SNE applicable other data as well, the proposed c-TSNE aims to only work well for scRNA-seq data. It is not even intended to apply to other similar data such as bulk RNA-seq data.

c-TSNE first assumes three important biological factors hierarchically contributing to cell diversity. The first is ‘which genes are expressed in single cell sequencing?’. Such a factor can be essential for discriminating two scRNA-seq samples because a large number of genes may not be expressed due to the dropout issue. The second is ‘what are the expression levels of the top-expressed genes in sequencing?’ Since the top-expressed genes can contribute to distinguishing cells more than the other genes, their impacts should be accounted for in sample similarity evaluation. The third is ‘what are the expression levels of whole genes?’. The third factor would play an important role in distinguishing cell diversity, provided two cells had the same or similar levels of contributions from the previous two factors.

c-TSNE then addresses the three biological factors by using appropriate and explainable distance metrics including Yule’s Y, low rank approximation with Chebyshev (L-Chebyshev), and fractional distance. The distance metrics aim to capture the differences between cells by analyzing the impacts from the gene expression status, the expression of the top-expressed genes, and the expression of whole genes jointly. The Yule metric detects the cell differences caused by that whether certain genes are expressed or not. It is employed to model scRNA-seq data by interpreting each zero entry as ‘gene not expressed’ and a non-zero entry as ‘gene expressed’. The L-Chebyshev metric is designed to capture impacts from those top expressed genes on the cell differences. The fractional distance metric is further adopted to evaluate the cell differences by considering the expression of the whole genes to avoid the bias from using the original Euclidean distance [9]. Finally, c-TSNE fuses the three metrics to build a more explainable and representative pairwise matrix according to input data sparsity. The fused pairwise distance matrix is expected to discriminate the cell similarities better for the fusion of the biologically meaningful distances. It is further employed to calculate the Gaussian distribution *P* in the input space and the student t-distribution *Q* in the embedding space before computing the final c-TSNE embedding [10].

c-TSNE is more explainable than the original t-SNE for its targeted customization. It interprets the diversity between cells fusing three biologically relevant ‘cell-driven’ distance metrics. The distance metrics, which aim to overcome the cell similarity evaluation bias of the Euclidean distance, are designed to consider the special characteristics of highdimensional sparse scRNA-seq data [9]. The experimental results show that scRNA-seq visualization and segregation with c-TSNE can not only much outperform the original t-SNE, but also state-of-the-art sophisticated peers such as SIMLR [8]. More importantly, it is easily understood and explainable than its peers from both scRNA-seq analysis and machine learning perspectives. To the best of our knowledge, it is the first explainable t-SNE for scRNA-seq analysis that integrates good interpretation with decent performance. Compared to other scRNA-seq clustering methods, the c-TSNE segregation can detect high-resolution subpopulations in more explainable ways. The robustness analysis shows that c-TSNE would demonstrate robustness to dropout and noise in clustering. Thus, it will inspire future explainable dimension reduction algorithm development in scRNA-seq analysis or even other biomedical data science fields.

## Results

The effectiveness and superiroity of c-TSNE can be validated by conducting single cell clustering besides straightforward visualization. The better the c-TSNE clustering quality, the more effective c-TSNE discovers the latent data structure of input scRNA-seq data. c-TSNE clustering consists of two steps. The first is to calculate the c-TSNE embedding of input single cell data. The second is to employ a clustering algorithm, say K-means, to cluster the embedding to discover the subpopulations. Although different clustering approaches can be employed, K-means is widely used for its popularity and simplicity though it needs to know the cluster number in prior [8]. Similarly, the other dimension reduction methods’ single cell clusterings follow the same scheme, i.e., it only needs to replace the c-TSNE embedding with the embeddings from the other dimension reduction methods such as t-SNE, UMAP or kernel principal component analysis (KPCA) [15–18].

### Peer methods

We compare the performance of c-TSNE in scRNA-seq clustering on benchmark datasets with four peer methods: t-SNE, UMAP, KPCA, and SIMLR [8–10]. They are either methods based on state-of-the-art dimension reduction techniques or widely accepted single cell clustering methods [8]. Unlike c-TSNE, the first three generally use the Euclidean distance to evaluate the cell similaries in dimension reduction. Both UMAP and KPCA demonstrate better performance than widely-used PCA (data not shown) according to our study. UMAP can be viewed as a method closely related to t-SNE because it can be somewhat viewed as a manifold learning method to fix some weakness of t-SNE (e.g., low convergence) [15–16]. KPCA is one of few methods to conduct dimension reduction in a high-dimensional Hilbert space by using kernel tricks [18]. We choose the cosine kernel *k*(*x*, *y*) = (*x* · *y*)/||*x*|| × ||*y*|| in KPCA for its automatic L2 normalizataion and good performance compared to the other popular kernels (e.g., Gaussian kernels) [18]. As a classic single cell clustering method, SIMLR seeks the learned optimal distance metric via multi-kernel learning to evaluate cell similarities in clustering [8].

### Benchmark single-cell RNA-seq datasets

We first use four widely used benchmark datasets to evaluate the performance c-TSNE clustering. Table 1 summarizes the basic information of the four datasets along with their sparsity values.

1. The Buettner dataset consisting of 182 cells across 3 classes was obtained from a controlled experiment which studied the effect of the cell cycle on the gene expression levels in individual mouse embryonic stem cells (mESCs) [19].
2. The Kolod dataset, which has 704 cells across 11 cell subpopulations, was obtained from a controlled experiment that studied the effect of the cell cycle on the gene expression level in individual mouse embryonic stem cells (mESCs) [18].
3. The Pollen dataset that consists of 249 cells across 11 cell populations including neural cells and blood cells (Pollen data set [20]). It was designed to test the utility of low-coverage single-cell RNA-seq in identifying distinct cell populations, and thus contained a mixture of diverse cell types: skin cells, pluripotent stem cells, blood cells, and neural cells.
4. The Usoskin dataset contains 622 neuronal cells from the mouse dorsal root ganglion across different 4 neuronal cell types, with an average of 1.14 million reads per cell [21]

**Table 1.** Bencgmark scRNA-seq data basic information

| Dataset | No. of Cells | No. of Genes | No. of classes | Sparsity (%) |
| --- | --- | --- | --- | --- |
| Buettner | 182 | 8989 | 3 | 37.9 |
| Kolod | 704 | 10685 | 3 | 27.9 |
| Pollen | 249 | 14805 | 11 | 51.0 |
| Usoskin | 622 | 17772 | 4 | 78.1 |

### Data sparsity

The sparsity is defined to measure the sparseness degree of an input scRNA-seq dataset X = {*x*_1_,*x*_2_,…*x_n_*}, 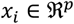, the sparsity of X is defined as the ratio between the number of zero entries over the total number of entries in the dataset: 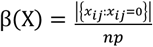. Data sparsity was rarely addressed in existing scRNA-seq studies, but it is a built-in characteristic of high-dimensional sparse scRNA-seq data because it is closely related to the dropout degrees in sequencing. Generally, the larger the sparsity, the more zeros appear in the dataset, and the more dropouts may happen in the transcriptome sampling procedure in sequencing. Since sparsity would affect the final fusion matrix in c-TSNE, and the Yule metric in c-TSNE is to address the data sparsity issue, it may deserve more attention in scRNA-seq analysis. Generally, the sparsity of a scRNA-seq dataset falls in the interval between 25% and 85% without imputation processing.

The cell-driven distances demonstrate good advantages over the default Euclidean distance in t-SNE visualization. Figure 1 shows the t-SNE visualizations under the Yule, L-Chebyshev, and Euclidean distance (default) across the four datasets, in which each point is colored by its cluster label in the corresponding dataset. It is interesting to see the t-SNE visualizations are more meaningful and representative under the two cell-driven metrics: t-SNE with the Yule and L-chebyshev metrics improve visualization interpretation by discovering more informative clustering structures. It clearly shows the well-separated three cluster separations for the Buettner and Kolod data. On the other hand, t-SNE with the Euclidean distance make cells jammed or even wired together and cannot indicate their cluster structures well. The default metric obviously makes the t-SNE visualizations less explainable because they cannot display the correct clusters due to its poor discriminability for high-dimensional sparse data. Similarly, t-SNE under the cell-driven metrics continue to demonstrate their advantages on the Usoskin and Pollen datasets by producing more orthogonal and separable clusters in visualization. The fractional distance also achieves leading advantages over the Euclidean distance in visualizations that can be found in the supplemental materials.

**Fig 1.**
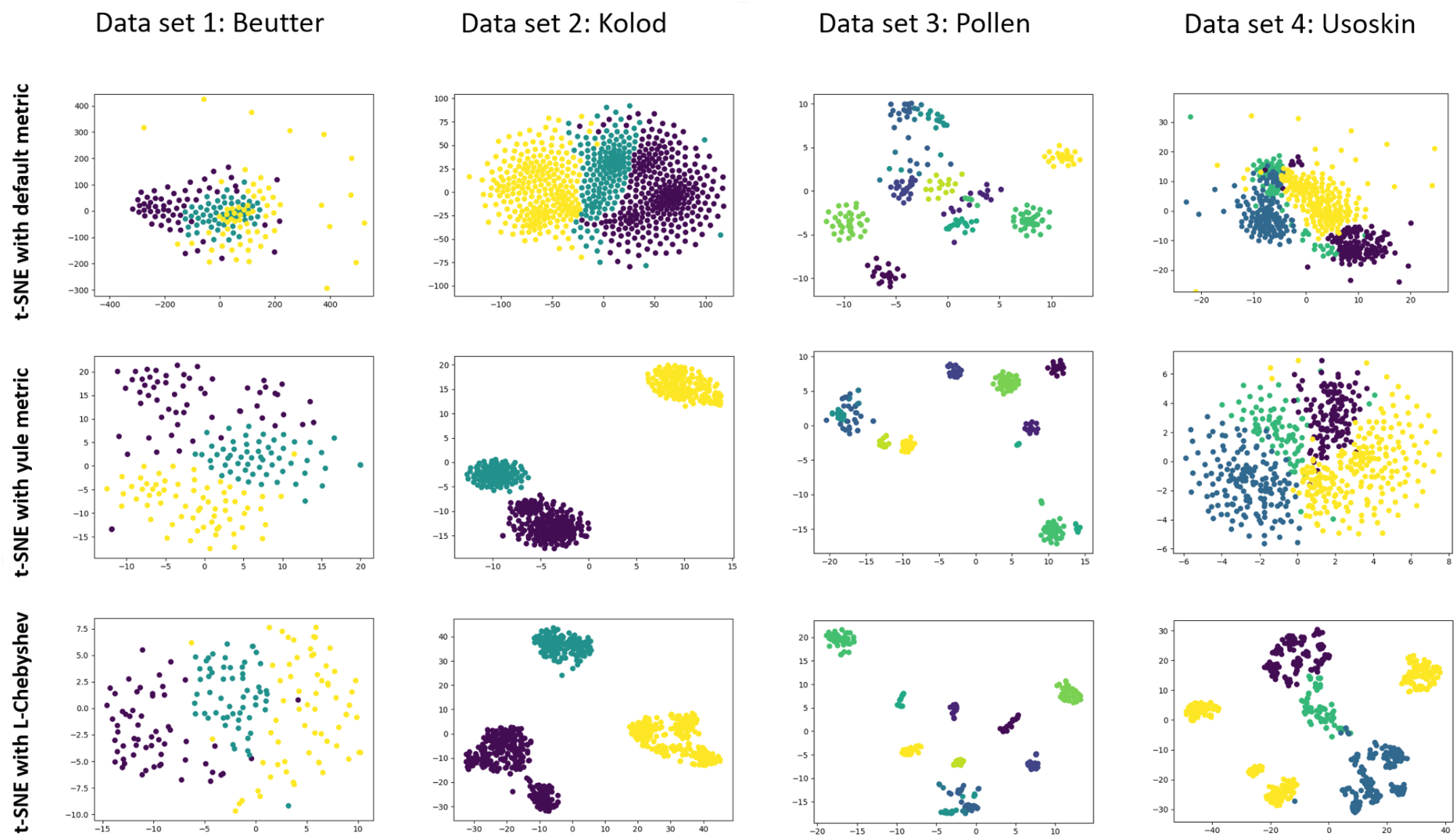
The t-SNE visualization comparisons under the Yule, L-Chebshev, and Euclidean distances (default) on the four benchmark datasets that have 3,3,11, and 4 clusters. The Yule and L-Chebshev distances demonstrate more meaningful separations for different subpopulations than the Euclidean distance for their more representative embeddings.

The t-SNE clustering results confirm the t-SNE visualization advantage under the cell-driven metrics over the default metric. The t-SNE clustering here means clustering different t-SNE embeddings under different distance metrics. The effectivness of clustering is evaluated by using a widely used clustering evalaution metric: normalized mutual information (NMI) that is a ratio between 0 and 1. The better the clustering quality, the closer the NMI value to 1. More details about NMI can be found in the Method section.

Table 2 compares the corresponding normalized mutual information (NMI) values of t-SNE clustering under the cell-driven and default metrics. It echoes the previous visualization results and suggests that the cell-driven metrics not only show high quality clustering performance for their higher NMIs, but make the clustering results more explainable. The more targeted distance metric choice, the better cell similarity discrimination, the more explainable t-SNE visualization, and the better cell segregation. It also shows that the widely used Euclidean distance has the worst performance for all datasets. For example, the NMI reaches 1.0 under the Yule and L-chebyshev for the Kolod data, but only 0.47 under the default Euclidean. Similarly, the NMI reaches 0.79 for the Buettner data under the fractional distance, but only 0.18 under the Euclidean. The results indicates that the cell-driven metrics can unveil the cell differences in an explainable and effective way as well as confirm the previous result that the Euclidean distance can be biased for high-dimensional data [9].

**Table 2.** The NMIs of t-SNE clustering under different metrics

| *Datasets* | *Yule* | *L-chebyshev* | *Euclidean* | *Fractional* |
| --- | --- | --- | --- | --- |
| *Buettner* | 0.73 | 0.68 | 0.18 | 0.79 |
| *Kolod* | 1.0 | 1.0 | 0.47 | 0.68 |
| *Pollen* | 0.9 | 0.93 | 0.86 | 0.91 |
| *Usoskin* | 0.45 | 0.77 | 0.2 | 0.54 |

### c-TSNE visualization and segregation

c-TSNE visualizations are not only more meaningful than classic t-SNE visualization for its biologically explainable distance metric usage, but also more representative than t-SNE visualizations on single cell-driven metrics. Figure 2 shows the embedding visualizations of c-TSNE in comparison with those of the classic t-SNE on the four datasets. The proposed c-TSNE produces more meaningful embeddings that generally show correct clustering than the classic t-SNE no matter with the sum or max fusion. Furthermore, c-TSNE demonstrates good advantages over t-SNE with individual cell-driven metrics. For example, the Buettner, Kolod, Pollen, and Usoskin datasets all show more orthogonal and better class bounds in visualization under c-TSNE than those under t-SNE with the individual cell-driven metrics. It suggests that the fusion procedure in c-TSNE contributes to improving the representative quality of the embeddings. It seems that max-fusion works well for the datasets with high sparsity. For example, the max-fusion produces the well-grouped four clusters for the Usoskin dataset with a 78.1% sparsity compared to the sum-fusion. On the other hand, the sum fusion seems to work well on the other datasets without high sparsity values.

**Fig 2.**
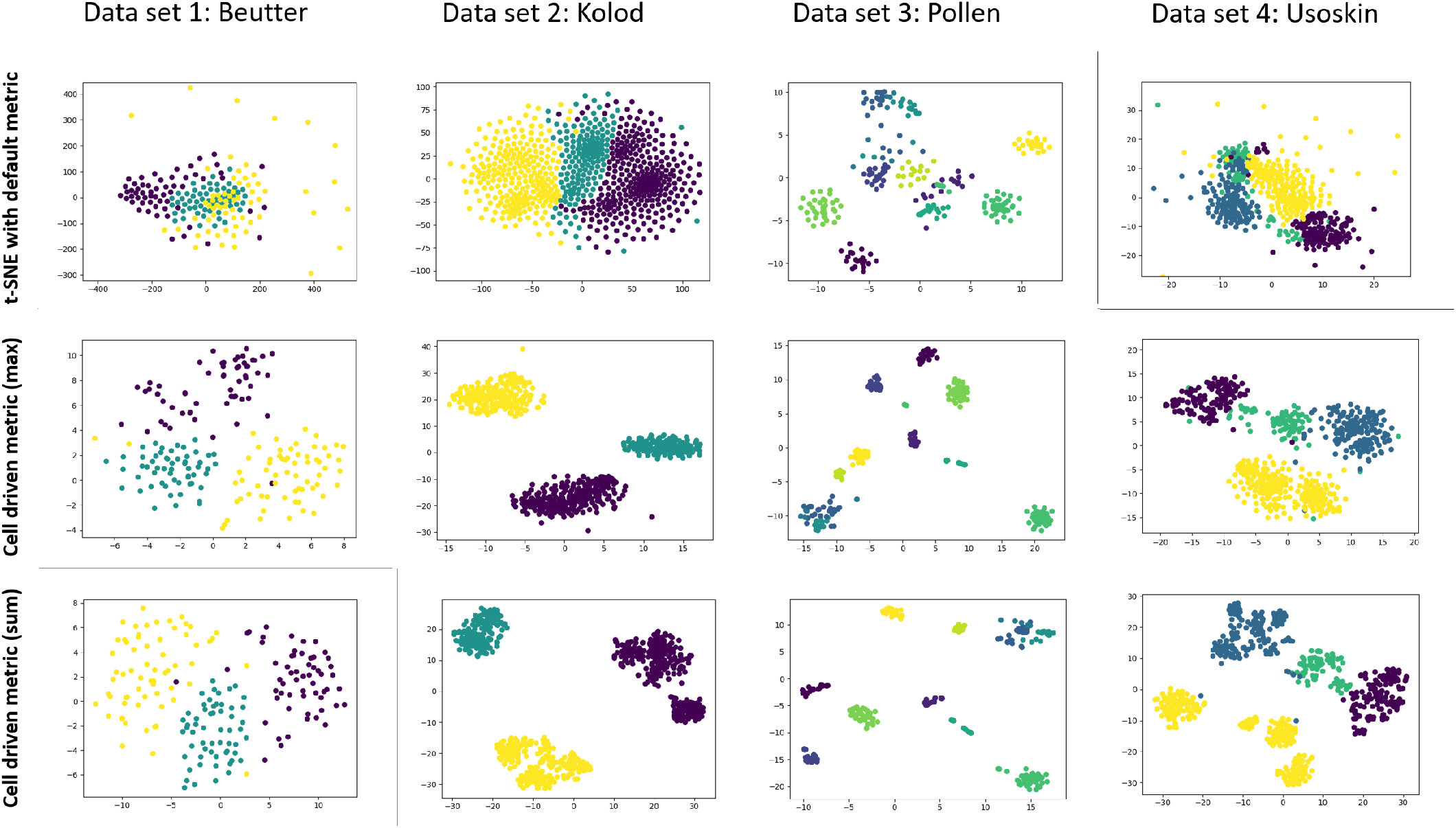
The comparison of the visualizations of c-TSNE and classic t-SNE with the default Euclidean metric on the four benchmark datasets that have 3, 3, 11, and 4 clusters respectively. c-TSNE creates more meaningful and separable clusters than the classic t-SNE in visualization for the four datasets no matter with the sum or max fusion as well as t-SNE under the individual cell-driven metrics.

c-TSNE clustering achieves stable and well-performed scRNA-seq segregations in a more explainable way. It clusters the c-TSNE embeddings by fusing different cell-driven metrics in the c-TSNE dimension reduction because the different biological factors represented by them may contribute to distinguishing the cell diversities jointly. Table 3 compares the NMIs from c-TSNE clustering with its peers: t-SNE, KPCA, and UMAP clustering and SIMLR, where the final NMIs from c-TSNE clustering are marked in bold. It shows that c-TSNE clustering shows leading and stable performance for all the four datasets in terms of NMIs. The NMIs from the t-SNE, KPCA, and UMAP clustering all demonstrate instable performance across the four datasets. Their good performance can be only achieved on an individual dataset but cannot be extended to the others. It suggests evaluating the cell similarity in t-SNE/UMAP with the default unexplainable Euclidean distance would lead to poor clustering performance. Only the performance of the complicated and expensive SIMLR method can compare with those of the c-TSNE clustering. Nevertheless, the c-TSNE clustering obviously outperforms SIMILR on the Kolod, Pollen, and Usoskin datasets in terms of the NMI values. It suggests that biologically explainable cell-driven distances in c-TSNE are more meaningful in single cell clustering than the ‘learned optimal distance metric’ in the SIMLR. Furthermore, c-TSNE clustering is more transparent, explainable, and implementation-friendly compared to the sophisticate SIMLR method [8].

**Table 3.**
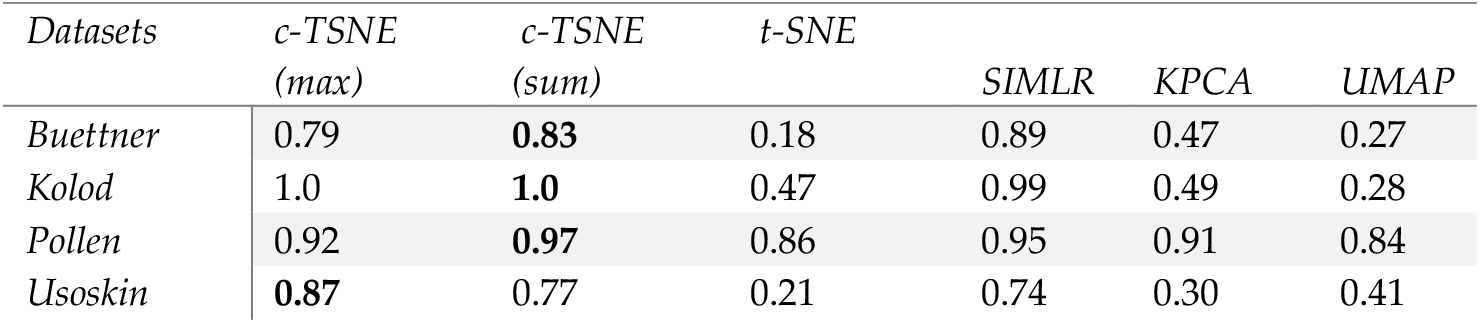
The performance comparisons of the c-TSNE clustering and its peers

### The entry-usage percentage under the max-fusion in c-TSNE

We examine the entry-usage percentage in the final fusion pairwise matrix of c-TSNE under the max-fusion (please go to Method for details). The max-fusion builds the final pairwise matrix for c-TSNE by selecting the maximum of the corresponding entries of the pairwise matrices generated by the three types of cell-driven distances. The entry-usage percentage is a ratio between the number of entries appearing in the final fusion pairwise distance matrix and the number of total final fusion matrix entries. For example, given *n* samples, the entry-usage percentage for the Yule metric is defined as the percentage that the Yule pairwise matric entries appear in the final fusion matrix:

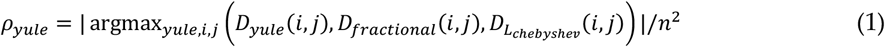

Examining the entry-usage percentage would unveil the max-fusion dynamics, provide insights about which cell-driven metric will have more contributions to the final fusion matrix in c-TSNE, and evaluate the role of sparsity in c-TSNE segregation.

We find that data sparsity can an important role in the max-fusion. Table 4 summarizes the percentages of entry-usage of the three metrics. Interestingly, Kolod, the dataset with the lowest sparsity, has the most important contribution from the L-chebyshev metric (e.g., 41.5% entry-usage). It suggests the impact of the top-expressed genes on the final pairwise matrix is more important than that of whether some genes are expressed or not, when input data has a low sparsity. In contrast, Usoskin, the dataset with the largest sparsity, has the most important contribution from the Yule metric (e.g., 41.7% entry-usage). It implies the impact of whether the certain genes are expressed or not on the final pairwise matrix can be more important than those of the other two, when input data has a high sparsity. The results also provide good insights in max-fusion and sum-fusion selection in c-TSNE: it is recommended the max-fusion (sum-fusion) for the high (low) sparsity data. The c-TSNE clustering results in the Table 3 are consistent with this, i.e.,. the max-fusion achieves the best performance for the Usoskin data and the sum-fusion has the best or equivalent performance for those datasets with relatively low sparsity (e.g., Buettner).

**Table 4.** The percentage of entry usage in the fused distance matrix of c-TSNE

| Dataset | $\rho_{yule}$ | $\rho_{fractional}$ | $\rho_{Lchebyshev}$ |
| --- | --- | --- | --- |
| Buettner | 34.3 | 34.8 | 30.9 |
| Kolod | 25.6 | 32.9 | 41.5 |
| Pollen | 26.7 | 38.2 | 35.1 |
| Usoskin | 41.7 | 23.6 | 34.7 |

### The robustness tests of c-TSNE clustering

It is necessary to examine the robustness of c-TSNE clustering to validate how it is resistant to dropouts and noise. This is rarely mentioned in single cell segregation in the previous literature, but it can be essential for single cell clustering because the dropouts and noise can be viewed as built-in components of scRNA-seq data. Only the methods passing the robustness tests would own good generalization in clustering. On the other hand, because the c-TSNE clustering mainly relies on the embeddings produced from c-TSNE, it is actually the robustness test of the proposed c-TSNE. Since the dropouts are a built-in nature of scRNA-seq data, it is highly likely that the sparsity of a given dataset would vary much from the original one provided the sequencing were conducted the second time. Thus, it needs to answer how robust the proposed c-TSNE will under different dropout events.

Moreover, noise can be involved in the gene expression quantification of scRNA-seq data from different sources that range from experimental design, human error, or even unknown system or biological complexity issues [6,13,14]. Therefore, it also needs to answer how robust c-TSNE will behave under the involvement of random noise. These queries are rarely investigated in the previous low-dimensional methods-based scRNA-seq clustering where each dataset is assumed as the final deterministic one. However, answering these questions will help us know whether c-TSNE clustering can be applied to solve generic single cell data segregation rather than only provide a case study.

To test the robustness of c-TSNE clustering, we conduct the robustness test for each dataset to evaluate the impacts of dropouts and noise involvement on the c-TSNE clustering in comparison with the classic t-SNE clustering. In the robustness test for dropouts, we view each dataset as a population and randomly drop various data fractions that range from 5% to 50% of the original data to analyze the performance of c-TSNE clustering by observing the changes of their NMIs.

Figure 3 illustrates that the robustness test results on dropouts for c-TSNE and t-SNE clustering on the four datasets. It shows that c-TSNE clustering always keeps the leading advantage over t-SNE clustering in terms of NMIs in a large margin under various dropout fractions. The NMIs of c-TSNE clustering are much larger than those of the t-SNE clustering for all datasets under different dropout fractions stably. For example, the NMIs of c-TSNE clustering have the least changes under different levels of dropouts on the Kolod dataset. In contrast, the NMIs of t-SNE clustering show large oscillations under dropouts, which is especially true for the Usoskin dataset with the largest sparsity. Similarly, it also suggests the robustness of c-TSNE to dropouts, i.e., c-TSNE embeddings are more stable with respect to different levels of dropouts than the original t-SNE embeddings. It further implies that c-TSNE clustering can achieve robust performance with good generalization.

**Figure 3.**
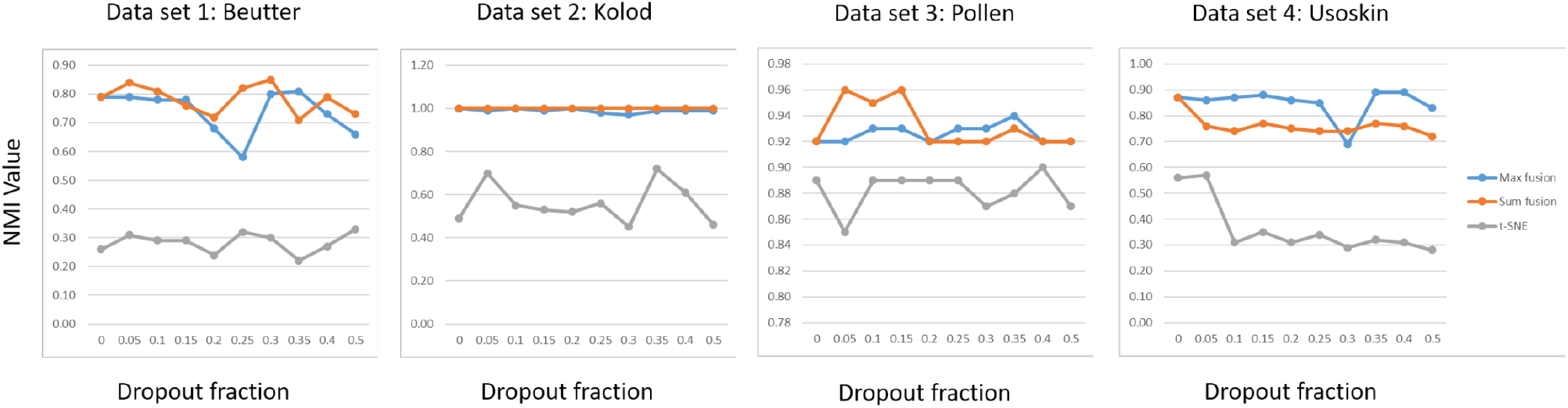
The robustness test of dropouts on the c-TSNE segregation and classic t-SNE clustering. The c-TSNE clustering keeps the leading performance over t-SNE clustering in a large margin stably for all the four datasets under different dropout fractions. On the other hand, the NMIs of the t-SNE clustering show a large level of oscillations under dropouts, which is especially true for the Usoskin data that has the largest sparsity. The NMIs of the c-TSNE clustering have the least changes under different levels of dropouts on the Kolod dataset that has the least sparsity.

Figure 4 illustrates the robustness test of random noise involvement in c-TSNE clustering in comparison with t-SNE clustering in terms of NMIs. It shows that c-TSNE clustering holds consistently leading advantages over t-SNE clustering under random noise with different levels of variances. For example, the maximum NMI of t-SNE clustering of the Buettner dataset is only 0.30 but the minimum NMI of c-TSNE clustering is about 0.70 under different levels of noise involvements. The similar patterns can be found for the other datasets also. To conduct the robustness test with respect to random noise involvement, we add independent zero-mean Gaussian noise with different levels of variances by following *x_ij_* = *x_ij_* + *N*(0, *σ_ij_*), 0.1 ≤ *σ_ij_* < 0.7, to input single cell RNA-seq data.

**Figure 4.**
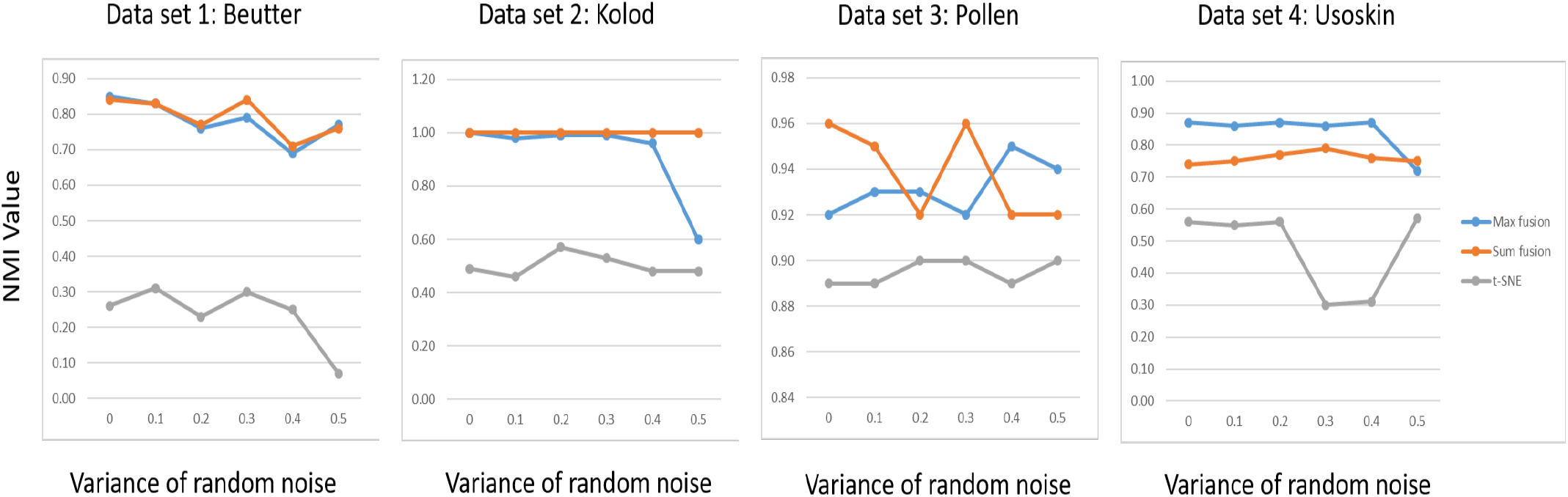
The robustness test of noise involvement on c-TSNE clustering and t-SNE clustering. The c-TSNE clustering demonstrates consistent leading advantages over the t-SNE clustering under different variances of random noise. The c-TSNE clustering seems to be more robust to noise involvement under the sum-fusion than the max-fusion.

We find no matter how the variance of random noise increases, c-TSNE clustering always leads t-SNE clustering by a large margin. This is probably because the cell-driven metric fusion weakens the impacts of the noise on clustering. Furthermore, c-TSNE clustering seems to be more robust to noise involvement under the sum-fusion than under the max-fusion generally. However, c-TSNE clustering with the max-fusion outperforms the sum-fusion on the Usoskin dataset that has the highest sparsity. It concurs that the c-TSNE with the max-fusion would be the better choice in cell segregation for those with high sparsity. In summary, we demonstrate the robustness of c-TSNE clustering with respect to dropouts and noise, suggesting it would perform well when applied to the other scRNA-seq data clustering.

### c-TSNE clustering effectiveness validation on additional datasets

We include two additional benchmark scRNA-seq datasets: Goolam and Patel to further verify the superiority of c-TSNE clustering [24–25]. Table 5 illustrates the basic information of the two datasets, both of which have relatively high sparsity. The Goolam dataset contains 124 blastomeres from 5 different stages of mouse embryos. In the Goolam dataset, transcriptomes were determined for all blastomeres of 28 embryos at the 2-cell, 4-cell and 8-cell stages, and individual cells taken from 16- and 32-cell stage embryos [25]. The Patel dataset consists of 430 cells from five heterogeneous primary glioblastomas according to their transcriptional expressions [24].

**Table 5.** Additional two benchmark datasets information

| Dataset | No. of Cells | No. of Genes | No. of classes | Sparsity (%) |
| --- | --- | --- | --- | --- |
| Patel | 543 | 5947 | 5 | 53.3 |
| Goolam | 124 | 41480 | 5 | 68.6 |

c-TSNE maintains the advantage over its peer methods in visualization and following clustering. Figure 5 compares the c-TSNE visualizations for the Patel data with it peers. The peer methods include t-SNE under the Yule, Fractional, and L-Chebyshev metrics, KPCA, UMAP, and the classic t-SNE. It seems that the visualization from c-TSNE with the sum-fusion achieve the best performance among all the methods, i.e., the five clusters are well separated compared to the other methods. The weight setting in the sum-fusion does not take the default uniform setting. Instead, we assign more weights to the Yule metric in the fusion because of the Patel dataset’s relatively high sparsity: *w_yule_* = 0.8, *w_fractional_* = 0.1, *w_L_chebyshev__* = 0.1. It is noted that c-TSNE with the max-fusion and t-SNE under the Yule metric can achieve almost equivalent visualizations for their good cluster separations. Table 6 shows their corresponding NMIs are 0.89, 0.87, and 0.86, respectively. This suggests that, for a high-sparsity dataset, whether certain genes are expressed or not can provide the most important information in data clustering compared to the other factors. It also explains why t-SNE under the Yule metric achieves the second-best clustering performance in terms of NMIs.

**Fig 5.**
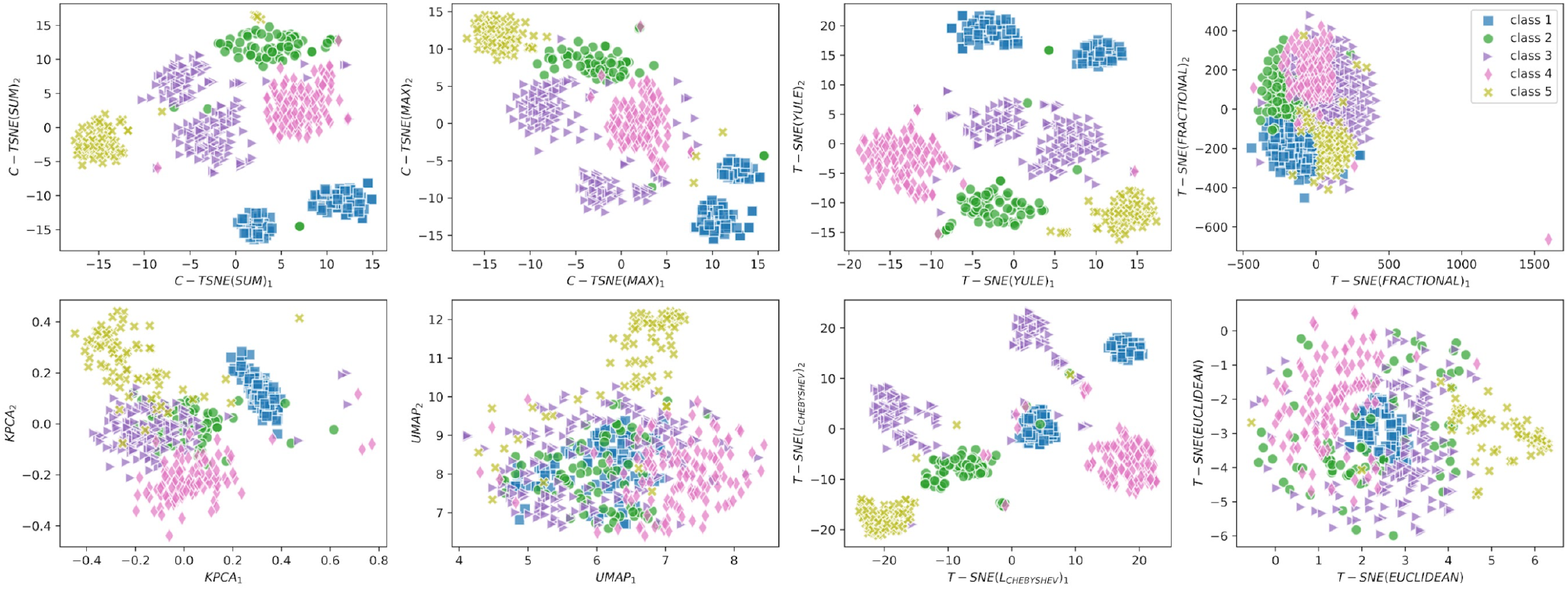
The comparisons of the c-TSNE visualizations with its peers including t-SNE visualizations with the individual cell-driven and Euclidean metrics, KPCA and UMAP visualizations for the Patel dataset. c-TSNE with the sum-fusion and max-fusion as well as t-SNE with the Yule metric achieve almost equivalent visualizations for their good cluster separations for this high-sparsity dataset. Both t-SNE and UMAP with the Euclidean metric have the worst cluster separations among all the methods. KPCA and t-SNE with the L-Chebyshev and fractional metrics all have decent separations though not the best ones.

**Table 6.**
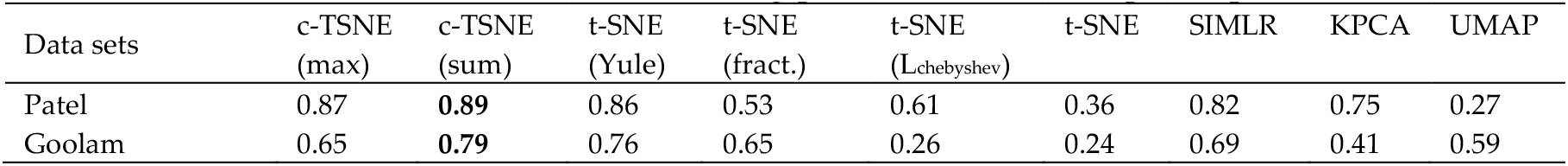
The cell-driven t-SNE clustering performance and its peers’ performance

| Data sets | c-TSNE<br>(max) | c-TSNE<br>(sum) | t-SNE<br>(Yule) | t-SNE<br>(fract.) | t-SNE<br>(Lchebyshev) | t-SNE | SIMLR | KPCA | UMAP |
| --- | --- | --- | --- | --- | --- | --- | --- | --- | --- |
| Patel | 0.87 | **0.89** | 0.86 | 0.53 | 0.61 | 0.36 | 0.82 | 0.75 | 0.27 |
| Goolam | 0.65 | **0.79** | 0.76 | 0.65 | 0.26 | 0.24 | 0.69 | 0.41 | 0.59 |

On the other hand, both UMAP and t-SNE under the Euclidean distance have quite poor performance, in which different groups of cells do not have clear boundaries (Fig 5). Compared to their t-SNE peers under the individual cell-driven metrics, the UMAP and t-SNE embeddings have the relatively small value ranges. For example, the ranges of the UMAP_1_ and UMAP_2_ embedding values fall in the intervals [4, 8.7] and [6.3, 12.5] and the ranges of the t-SNE_1_ and t-SNE_2_ embedding values fall in [-1, 6.7] and [-6.2, 0.9]. However, the ranges of the t-SNE_1_ and t-SNE_2_ embedding under the Yule metrics fall in much larger intervals: [-20, 19] and [-11,24]. The limited ranges of the embedding values limit t-SNE and UMAP to extract more meaningful latent data intrinsic structures as the others. On the other hand, it validates the previous result about the bias and limitation of using the Euclidean distance to evaluate high-dimensional data distance [9]. Furthermore, both KPCA and t-SNE with the L-Chebyschev metric achieve relatively decent separations for the dataset with NMIs 0.75 and 0.61 respectively, but the decent performance may not be consistent with all datasets from the previous results.

c-TSNE continues demonstrating its superiority to its peers on the Goolam dataset. Figure 6 compares c-TSNE visualizations for the Goolam data with it peers. c-TSNE with the sum-fusion demonstrates the obvious advantage in visualization for its almost well separated subgroups. The corresponding c-TSNE clustering achieves NMI 0.79, which is slightly higher than the NMI 0.76 achieved under t-SNE with the Yule metric. It indicates that the Yule metric can be more important than the other metric for the Goolam dataset with the 68.6% sparsity. Like other previous cases, t-SNE with the default Euclidean metric has still the worst visualization and clustering performance. t-SNE with the L-Chebyschev metric has a poor performance on this dataset though it has a decent performance for the Patel dataset. UMAP, KPCA, and t-SNE with the fractional metric achieve relatively fair cluster separations in their visualizations, although they may not separate all groups well for the Goolam dataset.

**Fig 6.**
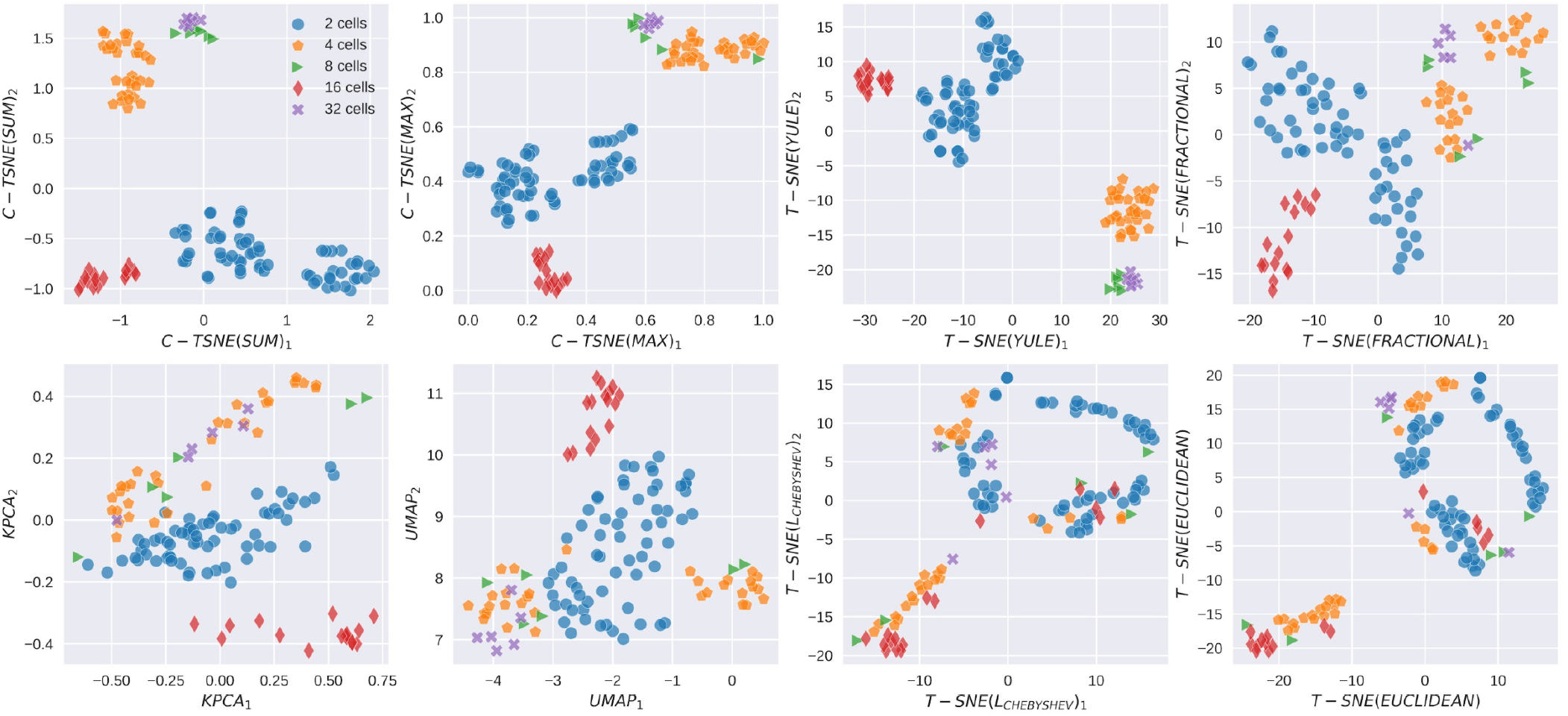
The comparisons of the c-TSNE visualizations with its peers: t-SNE with the three cell-driven and default metrics, KPCA, and UMAP visualizations for the Goolam dataset. c-TSNE with the sum-fusion/max-fusion and t-SNE with the Yule metric achieve almost the same level of cluster separations in their visualizations. The t-SNE with the default Euclidean distance and L-Chebshev metric both have the worst level of cluster separations. But KPCA, UMAP, and t-SNE with the fractional metric all have relatively fair cluster separations, especially the t-SNE with the fractional metric.

Table 6 compares c-TSNE clustering and its peers for the Patel and Goolam datasets, in which c-TSNE performance is marked in bold. Like before, c-TSNE clustering shows the best performance and stability among all methods. We notice that the Yule metric seems to outperform other metrics in clustering because of the relatively high sparsity of input datasets. In fact, almost all the cell-driven metrics demonstrate the obviously leading advantages over the default Euclidean distance under t-SNE clustering. For example, t-SNE with the Yule, fractional, and L-Chebshev metrics achieve 0.86, 0.53, and 0.61 NMI values in the Goolam data clustering, and 0.76, 0.65, and 0.26 in the Patel data clustering respectively. The best results from the Yule metric are even better than those from the complicate SIMLR clustering method [8]. On the other hand, KPCA and UMAP fail to achieve consistently good performance. For example, UMAP achieves a fair level NMI on the Goolam data, but it provides a very poor clustering with NMI 0.27. In summary, c-TSNE demonstrates its stable and good performance in visualization and clustering compared to almost all its peers.

To the best of our knowledge, c-TSNE clustering achieves the best performance or at least equivalent performance for all the six datasets in this study in an explainable and efficient way [26–27]. t-SNE with the default Euclidean distance has the worst performance for almost all datasets. It suggests the popularity using t-SNE with the default Euclidean distance as a visualization and clustering tool in scRNA-seq analysis may not be a good choice for its poor performance and lack of good interpretation in medicine and biology. The other peers all demonstrate instability in achieving good cell segreations. They are unable achieve consistently decent clustering performance for different datasets. For example, KPCA achieves 0.91 NMI for the Pollen data but only 0.41 NMI for the Goolam data. On the other hand, it suggests highly nonlinear and complexity of scRNA-seq data itself as well as the effectiveness and repeatability of c-TSNE.

## Discussion

c-TSNE provides an explainable visualization and clustering tool for scRNA-seq analysis. Unlike traditional t-SNE, it employs the cell-driven metrics to distinguish the cell differences in the low-dimensional embedding space. The Yule, fractional, and L-chebyshve distance metrics model the most relevant biological factors in scRNA-seq cell differences. The fused distance matrix from the metrics models the cell similarity more representative, biologically meaningful, and easily understood and explainable.

Moreover, c-TSNE somewhat breaks the general myth about the trading-off between machine learning efficiency and explainability [11]. It strongly demonstrates that enhancing the interpretability of a machine learning model (e.g., t-SNE) can contribute to its efficiency as well. In out context, it means to improve clustering performance and robustness to dropouts and noise. More importantly, our method stays explainable by avoiding multi-kernel learning or possible feature selection tuning that makes single cell segregation ambiguous or even a black-box procedure [8,26]. By combining different cell diversities revealed by different interpretable metrics, the proposed cell-driven t-SNE also shed light on the still unknown mechanism of cellular differentiation from an explainable machine learning perspective [3,11].

Unlike other existing scRNA-seq clustering methods, the proposed c-TSNE clustering conducts segreation from the raw scRNA-seq data rather than normalized one by viewing the raw data as the final data [28]. It views the dropout issue as a normal procedure in sampling the whole transcriptome. It considers the sparsity as an important parameter in deciding the final pairwise matrix fusion in c-TSNE. It is also one important reason we choose the raw data rather than the normalized data or data after imputation because data may lose its original sparsity after normalization or imputation [14]. To the best of our knowledge, it is the first single cell segregation method considering the sparseness degree of input data. That the results of c-TSNE clustering are superior to its peers suggests the effectiveness of using the raw single cell RNA-seq data for the sake of better single cell clustering. It may suggest imputation processing may not be an essential one for determining the cell-to-cell differences. The recent work reported that most imputation methods had no impact on scRNA-seq analysis compared to non-imputed data [29]. It would be interesting for us to evaluate the role of normalization in scRNA-seq clustering by employing normalized data using the widely used normalization methods (e.g., SCron) in c-TSNE from a clustering model selection perspective [12,26,29].

Furthermore, the proposed method does not rely on feature selection to determinate single cell data dimension reduction and following segregation because we believe feature selection may not be able to provide a good interpretation. In other words, we invite all transcripts involved in the single cell segregation because we believe the whole transcriptome will determine the cell segregation rather than few important genes selected from some feature selection methods that may lack good interpretation. Since previous results reported that appropriate feature selection may contribute to scRNA-seq segregation, it would be worthwhile to compare c-TSNE clustering with the whole transcriptome and interpretable feature selection [26,29].

### c-TSNE Speedup

The weakness of the proposed cell-driven t-SNE (cTSNE) can be its high complexity though it is still implementable because most scRNA-seq datasets have their sample size <1000. However, it is desirable to decrease the complexity of c-TSNE so that it can handle datasets with a large number of cells, especially because of the increasing number of cells assayed per experiment in scRNA-seq [30,31].

We can optimize the existing complexity of the proposed c-TSNE to *O*(*n*^2^ + *nlogn*) with the following two speedup techniques. The first is to use less expensive method to conduct normalization for the three different pairwise distance matrices before fusion. Without the normalization procedure for three different distance matrices, c-TSNE has almost the same complexity as the classic t-SNE: *O*(*nlogn*). But the eigenvalue decomposition involved in the distance matrix normalization generally take *O*(*n*^3^) in practice. It is somewhat expensive to conduct such normalization by doing an eigenvalue decomposition though it does achieve good performance [32]. Using alterative less expensive normalization scaling factor *trace*(*D*)/*n* would greatly decrease the complexity cost because the trace is the sum of the eigenvalues, but it can be calculated by the sum of all. Thus, the complexity of cell-driven t-SNE should be decreased by using this technique.

The second is to use truncated SVD to calculate the L-chebyshev distance metric. The L-chebyshev distance that needs an SVD-based reconstruction may take more costs because SVD complexity can be *O*(*n*^3^). However, such a weakness can be fixed by using truncated SVD that has *O*(*n*^2^) [32]. Thus, the optimized cell-driven t-SNE is an implementable-favor method for its *O*(*n*^2^ + *nlogn*) complexity and will work well for possible datasets with a large number of cells.

## Conclusion

We provide an explainable t-SNE: cell-driven t-SNE (c-TSNE) for scRNA-seq data visualization and clustering. Its explainability lies in the fact it is a customized dimension reduction method designed only for high-dimensional sparse scRNA-seq data. The more explainable and biologically meaningful distance metrics are employed in c-TSNE to discriminate the cell differences rather than use the default Euclidean distance that may bring biased or even meaningless results for high-dimensional scRNA-seq data. It has no intention to generalize c-TSNE to other datasets such as bulk RNA-seq data or even normalized or imputed scRNA-seq data. The proposed explainable t-SNE outperforms classic t-SNE and its other peers in meaningful visualization and segregation in an easily understood and interpretable way. The poor performance from the classic t-SNE in visualization and clustering may suggest that t-SNE can be easily misused in the existing scRNA-seq analysis for its popularity. On the other hand, the poor performance of t-SNE highlights the importance of developing explainable machine learning methods in scRNA-seq analysis. The customized machine learning methods should be designed from an explainable perspective to enhance efficiency in each application domain, especially because most machine learning methods employed in biomedical data science fields are methods migrated from the other fields [33].

We recommend c-TSNE under the sum-fusion for most input data for its good performance and flexibility for different datasets. More future work may be needed to provide a more rigorous but easily implemented weight selection method though the existing one works. We observe the power of using the Yule metric in discriminating single cells besides the fractional and L_chebyshve_ metrics. It is possible to employ the Yule metric alone besides the max-fusion for some datasets with very high sparsity for the sake of efficiency according to the existing experimental results. However, not all high-sparsity datasets can achieve the best performance in visualization and following clustering. For example, the Usoskin dataset has the highest sparsity (78.1%), but it only achieves 0.45 NMI under the Yule metric alone. It is a possible to figure out how to set the weights for the Yule metric wisely in the existing sum-fusion. On the other hand, another Yule-similar but a more explainable pseudo-metric needs to be built from a nonlinear optimization approach to reflect the true cell similarity for the high-sparsity datasets [34].

Besides comparing c-TSNE clustering with other state-of-the-art peers such as SOAP [35], we plan to apply c-TSNE to single cell classification to further validate its superiority and possible enhancement as well as compare it with the existing cell classification methods [36–37]. For example, it is possible to make the existing distance fusion more accurate and adaptive by considering the possible impacts from other factors (e.g., data entropy) in the specific classification procedure, besides the existing data sparsity because it is likely that other latent data characteristics (e.g., data entropy) can also affect fusion besides sparsity [38].

The existing clustering procedure for the embedding from cell-driven t-SNE and its peers is K-means. It has the advantage of simplicity and efficiency because of its explainability, but it requires the input embeddings to be convex, which may not be guaranteed each time for those c-TSNE, t-SNE, or UMAP embeddings obtained from dimension reduction. Also, K-means needs to know the number of clusters in advance to have a good estimation. It may be desirable to try different clustering methods such as DBSCAN that can overcome the weakness of the K-means in the clustering quality evaluation [39]. At the same time, we are also interested in investigating to extend the proposed c-TSNE to its quantum version to exploit its quantum advantage to handle larger and more complicated scRNA-seq datasets in a fast and accurate way [40].

## Method

Before describing the proposed cell-driven t-SNE (c-TSNE), we introduce cell-driven diversity distance metrics as follows. The metrics include the Yule metric (Yule’Y), low-rank approximation Chebyshev (L-Chebyshev) metric, and fractional distance metric [41–44]. The cell-driven metrics are used to evaluate the impacts from the three biologically meaningul factors on cell differences: ‘which genes are expressed in single cell sequencing’, ‘ the expression levels of the top-expressed genes’, and ‘the expression levels of whole genes’ respectively.

### The cell difference under the Yule metric

The Yule metric, namely *Yule’s Y*, developed by George Udny Yule in 1912, is a measure of association between two binary variables and known as the coefficient of colligation [41]. To measure the cell differences under the Yule’s metric, we first map each scRNA-seq sample *x* = (*x*_1_,*x*_2_,…,*x_N_*)*^t^*, to its binary vector 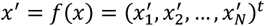 by using the mapping function 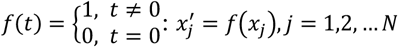. The binarization preprocess maps all zeros in a scRNA-seq sample to 0 (‘not expressed’) and all non-zero entries to 1 (‘expressed’), i.e., it creates a corresponding ‘binary map’ for each cell to model its gene expression status in sequencing.

Given the binary vectors of two scRNA-seq cells across *N* genes *U* = (*u*_1_,*u*_2_,*u*_3_,…,*u_N_*)^*t*^, *Z* = (*v*_1_,*v*_2_,*v*_3_…,*v_N_*)*^t^*, where *u_i_*, *v_i_* ∈ {0,1}, *i* ∈ {0,1,2,…,*N*}, then the cell difference between two cells caused by the statuses of gene expressed and not-expressed can be presented by the following Yule’s Y as follows,

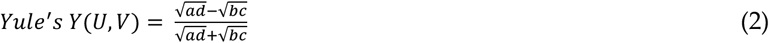

The parameter a is the frequency of both *v_i_* = 0 and *u_i_* = 0: *a* = |{*i*:*u_i_* = 0 ∧ *v_i_* = 1}|. Similarly, *b* = |{*i*:*u_i_* = 1∧ *v_i_* = 0}|, *c* = |{*i*:*u_i_* = 0 ∧ *v_i_* = 1}|, *d* = |{*i*:*u_i_* = 1 ∧ *v_i_* = 1}|. The range of Yule metric is between −1 and 1, where −1 and 1 reflect total negative (positive) association, whereas 0 reflects no association.

If the distance between two cells A and B under the Yule metric is 0.92, it means they are similar with respect to each other. Those genes expressed in the cell A will also express themselves in the cell B. Similarly, if the between two cells A and B under the Yule metric is −0.88, it means they are dissimilar with each other 88%. Those genes expressed in the cell A mostly will not express themselves in the cell B.

### The cell difference under low-rank approximation with Chebyshev metrics (L-Chebyshev)

The top-expressed genes in scRNA-seq data can contribute to a large portion of the whole data variances and play an important role in contributing to the cell differences. The top-expressed genes can be those differentially expressed with a large-fold expression change relevant to the other genes in one or more cells. Biologically, they may be important candidates in the pathway interacting with other genes mostly or even affecting other genes’ expressions. We design a low-rank approximation with Chebyshev metric (L-Chebyshev) to model the expression levels of the top-expressed genes as follows.

We first employ single value decomposition (SVD) to approximate the original data in the subspace spanned by the top singular vectors [42]. Given X = {*x*_1_,*x*_2_,…,*x_n_*}, 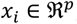, *p* ≫ *n* we have the SVD decomposition at the selected rank *l* < *p*

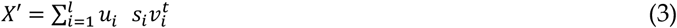

The selected approximation rank can be decided by 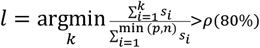. The approximated gene expression matrix 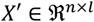 would include the expressions of the top-expressed genes and the contributions of those genes with low expressions are ignored in the SVD approximation. The maximum approximation rank *l* will be determined by checking the variance ratio 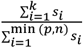 so that singular values of the selected approximation rank can hold at least 80% total variance ratio. As a result, each column vector in the matrix *X*’ is the approximated cell containing the expressions of the top-expressed genes corresponding to each original cell.

We then use the Chebyshev metric to capture the difference between two approximated cells to evaluate the impacts of top-expressed genes on the cell difference and such metric is named as the low-rank approximation with Chebyshev metric (L-Chebyshev). Given two cells *x*, 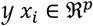 in *X*, the distance given by L-Chebyshev metric is calculated as follow:

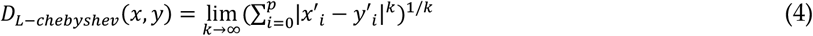

where the *x*’, 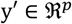 are the corresponding approximated cells of *x*, *y* in *X*’. The reason we use the Chebyshev distance rather than the default Euclidean distance is to avoid the bias from the Euclidean distance because the approximated cells are still high-dimensional data. In fact, the Chebyshev distance between two points is always less than the Euclidean distance for two approximated cells. Thus, it avoids the amplified Euclidean distances for the sake of more accurate evaluations of the impact of the expression of top-expressed genes on cell difference.

### The cell difference under fractional distance metrics

As we mentioned before, Aggarwal *et al*. mathematically proved that in the high dimensional space, the distance or similarity calculated by standard metrics, such as Euclidean metric, is likely to be meaningless or biased and named this phenomenon as ‘the curse of dimensionality.’ [9]. When the dimensionality becomes higher, the difference between the maximum and minimum distances, for certain data points, does not increase as fast as the minimum distance to other data points. When the dimensionality becomes extreme high, the ratio of the difference between the maximum and minimum distances to the minimum distances will be close to 0, which makes it hard to tell which data points are similar or dissimilar.

Inspired by Aggarwal et al’s work, we extend the normal Minkowski distance to a specific fractional distance metric to deal with this problem to evaluate the cell difference caused by all genes. Here, the fractional distance between the two single-cell samples *U*, 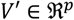 is defined as:

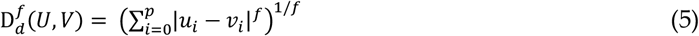

where f ∈ (0,1). Thus, we can augment the difference between the nearest point and the farthest point and give a better measurement of similarity in the high dimensional space. Technically, we can set any value for *f* in the interval, but a too small *f* value may bring some unnecessary oscillations in distance calculations for a relatively dense sample that with less zeros and another sample with more zeros. We empirically choose *f* = 1/4 in this study because it demonstrates a better sensitivity in detecting cell difference than the other values.

Although the fractional distance theoretically may not follow the triangle inequality for a distance metric for some very special points such as *x* = (0,0,…0)^*t*^, *y* = (1,0,…1)^t^, *z* = (0,0,…1)^t^, such exceptional cases are unlikely to happen for real scRNA-seq samples because there is no a real scRNA-seq sample with all zero expressions. Actually, the good performance from the fraction distance in clustering demonstrates that it would be an effective distance metric to discriminate cell difference better than the traditional measures practically. Thus, we can still treat it as a distance metric in evaluating the similarities between cells.

### Cell-driven distance fusion

The three different distance metrics applied to input scRNA-seq data generate three pairwise distance matrices to reflect the cell difference from the three different biological aspects. It needs to fuse the three matrixes to build a new pairwise distance matrix to model the cell difference more accurately, because the three biological factors contribute to the cell diversity jointly. A normalization process needs to apply to each matrix to standardize their scales so that they can be fused on the same page without losing their own characteristics. We find that standardizing each distance matrix to zero mean and one standard deviation will not be a good way because the distances may not be subject to a normal distribution, which is especially true for the distance matrix generated under the Yule and fractional distance metrics. Therefore, we adopt the approach to normalize each distance matrix by employing its largest eigenvalue as a scaling factor in normalization [16]. Given a distance matrix *D*, we conduct eigenvalue decomposition as follows,

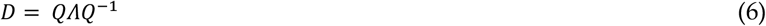

Then the normalized distance matrix 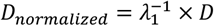, where *λ*_1_ = *eig_max_*(*D*) is the largest eigenvalue of the pairwise distance matrix *D* under a metric. It is noted that such a normalization factor scales each matrix well but can be expensive because the complexity of the eigenvalue decomposition can reach *O*(*n*^3^). But such a complexity should not be a concern in implementation because *n*, which represents the number of cells, will not be a number larger than 10^3^, for almost all scRNA-seq datasets.

After all pairwise distance matrices complete normalization, we fuse the three matrices to a new pairwise distance matrix for t-SNE. The ideal way is to assign exact weights to each matrix so that their impacts to the cell difference can be counted accurately. However, we lack enough prior knowledge to implement it because we do not know which factors will have more impact on the final cell diversity because of biological complexities, stochastic nature of sampling, dropout, and other artifacts. It is possible to do individual study for each dataset and their medical/biological background to determine a more appropriate fusing model for each dataset, but such a sophisticated fusing may let cell-driven t-SNE lose generalization for generic scRNA-seq data and increase algorithm complexity. Thus, we just take the following simple but effective fusions: max-fusion and sum-fusion described in the following two equations. The sum-fusion is recommended for most data for its generalization and good performance.

### Max-fusion & sum-fusion

Biologically, the max-fusion assumes the most important discriminated distance between two cells can come from any source no matter the top-expressed genes or others. It picks the *ρ_max_* value of corresponding entries from the three normalized distance matrices: *D_yule_*,*D_fractional_*, and *D_L_chebyshev_* as the corresponding entry of the fused matrix *D^f^*. The *ρ_max_* by default is the maximum function in our c-TSNE implementation.

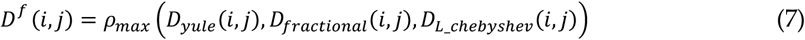

It is noted that *ρ_max_* can also be implemented as a percentile function to pick a high percentile (e.g., 80^th^ percentile) from the three pairwise matrix entries when the sparsity of the input dataset is high enough (e.g. 50%).

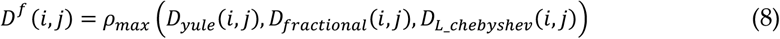

The summation fusion, i.e., sum-fusion, assumes the three sources contribute to the cell differences jointly in a weighted way:

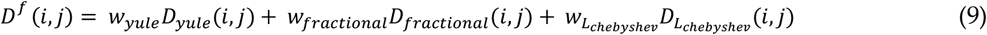

The weights *w_yule_*, *w_fractional_*, *w_L_chebyshev__* would not be determined in an accurate way without enough priori knowledge about data. The default weight selection would be 1/3 for each item under the situation. However, when input data has a high sparsity value (e.g., >0.5), it is recommended to adjust the weight of the Yule metric *w_yule_* to a large one (e.g., 0.8) according to data sparsity.

Generally, the sum-fusion is especially recommended for most input data for its generalization and good performance, especially for those with relatively small sparsity values, but max-fusion is recommended to handle the datasets with relatively large sparsity degrees. In some special cases, it is recommended to only take a single cell-driven metric such as the Yule metric for some datasets with a high degree of sparsity because gene expressed or not can be a more important factor in discriminating the cell difference compared to the other factors. More details can be found in the following algorithm.

### Cell-driven t-SNE (c-TSNE), an explainable t-SNE

The key difference between c-TSNE and original t-SNE lies in that c-TSNE calculates a biologically meaningful and cell-driven pairwise matrix *D^f^* fused by explainable distance matrices to evaluate cell similarity. We describe c-TSNE with more details by following the original t-SNE framework.

Given an input scRNA-seq dataset *X* = {*x*_1_,*x*_2_,…}, 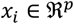, *P* ≫ *n*, c-TSNE calcualtes the corresponding low-dimensional embedding Y = {*y*_1_,*y*_2_,…*y_n_*}, 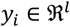, *n* ≫ *l*, generally *l* = 2, by minimizing the Kullback-Leibler (K-L) divergence between a Gaussian distribution *P* and a normalized Student’s t-distribution *Q*,

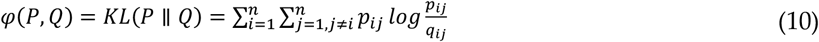

where *p_ij_* models the pairwise similarity between points *x_i_* and *x_j_* in the original high-dimensional manifold and *q_ij_* models the pairwise similarity of their corresponding low-dimensional embeddings: *y_i_* and *y_j_* [10]. 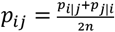 is defined as the average of two conditional probabilities, in which 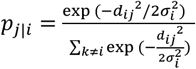, where *x_j_*, *x_k_* are the neighbors of *x_i_*, 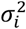 is the variance of all neighbor data points of *x_i_*, and *d_ij_*, = *D^f^* (*i*, *j*) is the fused distance between samples *x_i_* and *x_j_*. Similarly 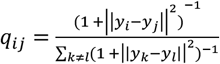. Due to the different input pairwise matrices, the distributions *P* and *Q* in the proposed c-TSNE will be totally different from those in the original t-SNE.

The c-TSNE embedding *y_j_*, is calculated by using the following gradient learning scheme,

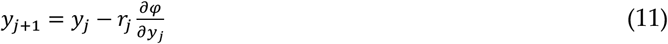

where 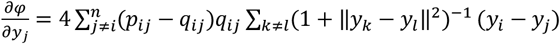, and *r_j_* is the learning rate, which is recommened to set as about 1/10 of the learning rate of the classic t-SNE. Although small learning rate might ‘condense’ the observations in small regions, we find the small learning rate will contrbute to finding better local optmimum under the cell-driven distances rather than got stuck in one local optimum easily though it takes more time [45].

It is noted that the embeddding from c-TSNE should be more representative and robust compared to the embedding from the original t-SNE because the fused distance matrix *D^f^* is more representative and robust compared to the one from the original t-SNE. The proposed c-TSNE still shows randomness as the classic t-SNE because of the nature of the non-unique solutions from the non-convex optimization [44]. However, c-TSNE tends to be more stable than the classic t-SNE under the change of the perplexity values.

The proposed c-TSNE has a higher complexity than the classic t-SNE: *O*(*n*^3^ + *nlogn*). It is due to the high complexities from distance matrix normalization and L-Chebyshev distance calculations, both of which need *O*(*n*^3^) complexity. However, since c-TSNE mainly handles high-dimensional scRNA-seq data, the number of *n* is mostly less than 1000. Therefore, c-TSNE is a technically workable algorithm though it appears as a high-complexity algorithm before going through complexity optimization strategies [44–45].

### c-TSNE segregation

c-TSNE segregation is to employ c-TSNE to calculate the low-dimensional embedding of input scRNA-seq data. Then a clustering algorithm is employed to cluster the c-TSNE embedding. As mentioned before, K-means is selected in this study for the conveience of comparisons because it is widely used in the existing literature for its simplicity and efficiency [8]. However, it can be easily replaced by other more complicate clustering algorithms such as DBSCAN or OPTICS [39,48].

The effectivness of clustering is evaluated by using classic clustering evalaution metrics such as normalized mutual information (NMI) [46–47]. The NMI, a ratio between 0 and 1, is defined as follows. Given two clustering results U and V for input data. then NMI is the ratio between the mutual information of *U* and *V* and the average entropy of *U* and *V*.

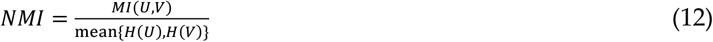

where 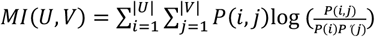, 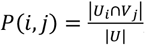, 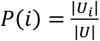, 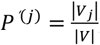, 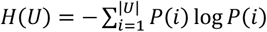, 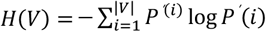. The larger the NMI indicates the better quality scRNA-seq data clustering [21]. Although other clustering measures are also available, the NMI demonstrates a good stability across different scRNA-seq datasets. Thus, we only choose NMI as the clustering quality index in this work.

The following algorithm describes the cell-driven t-SNE clustering, where c-TSNE is also described in a detailed way. The proposed c-TSNE seeks different fusion methods and even different weights according to the sparsity values of input data. When the input dataset has a sparsity β > the sparsity cutoff: which is set as 75% in this study, the max-fusion is employed to fuse the three pairwise distance matrices; when the sparsity falls in the interval [50%, 75%], more weights should be given to the weight of the pairwise distance matrix generated from the Yule metric.

The cell-driven t-SNE (c-TSNE) clustering has the complexity *O*(*n*^3^ + *nlogn* + *kn*), where *k* is the number of clusters and *n* is the number of observations. Compared to the complexity of the t-SNE-based clustering: *O*(*nlogn* + *kn*), it is slightly an expensive clustering algorithm, although it still can be done in a real-time because the scale of *n* < 10^3^.

#### Algorithm: Cell-driven t-SNE clustering

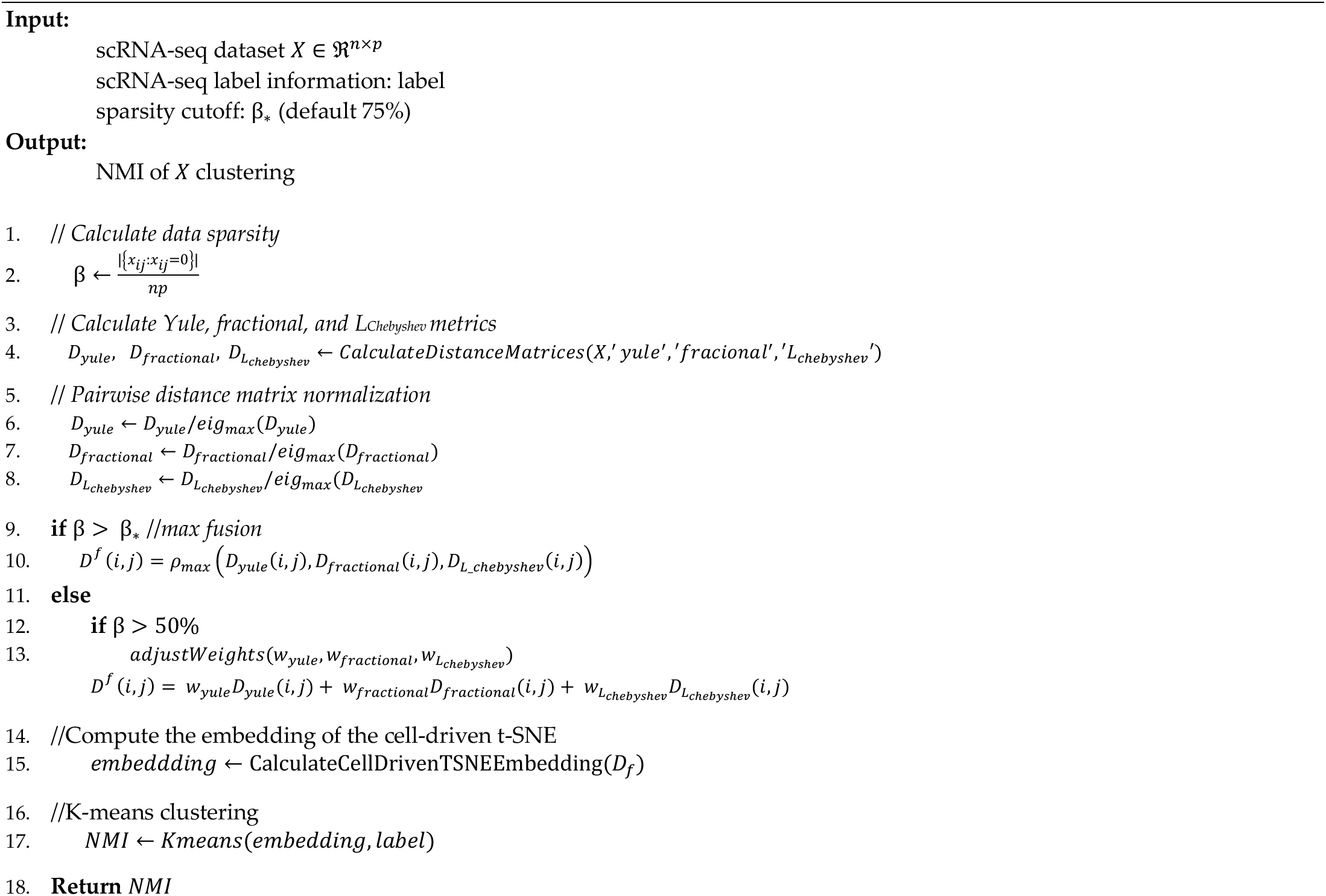

